# Image guided construction of a common coordinate framework for spatial transcriptome data

**DOI:** 10.1101/2023.11.14.565868

**Authors:** Peter Lais, Shawn Mishra, Kun Xiong, Gurinder S. Atwal, Yu Bai

**Affiliations:** Regeneron Pharmaceuticals, Inc., Tarrytown, NY; Georgia Institute of Technology

## Abstract

Spatial transcriptomics is a powerful technology for high-resolution mapping of gene expression in tissue samples, enabling a molecular level understanding of tissue architecture. The acquisition entails dissecting and profiling micron-thick tissue slices, with multiple slices often needed for a comprehensive study. However, the lack of a common coordinate framework (CCF) among slices, due to slicing and displacement variations, can hinder data analysis, making data comparison and integration challenging, and potentially compromising analysis accuracy. Here we present a deep learning algorithm STaCker that unifies the coordinates of transcriptomic slices via an image registration process. STaCker derives a composite image representation by integrating tissue image and gene expressions that are transformed to be resilient to noise and batch effects. Trained exclusively on diverse synthetic data, STaCker overcomes the training data scarcity and is applicable to any tissue type. Its performance on various benchmarking datasets shows a significant increase in spatial concordance in aligned slices, surpassing existing methods. STaCker also successfully harmonizes multiple real spatial transcriptome datasets. These results indicate that STaCker is a valuable computational tool for constructing a CCF with spatial transcriptome data.

## Introduction

Spatial transcriptomics is a cutting-edge OMICS technology that enables characterizing transcriptome-wise gene expressions at spatially resolved locations in a tissue section (1,2). The data acquisition involves dissecting a thin slice (5-20 micrometers) from a tissue block for subsequent molecular analyses. Multiple slices are often profiled to ensure data reliability. However, each slice is analyzed separately due to variations in cutting and displacement or by sample attrition. Integrating multiple slices into a common coordinate framework (CCF) is critical to enhance resolution, detect spatial patterns, and build a three-dimensional perspective of the tissue microenvironment from two-dimensional profiles.

Despite the importance of the CCF, existing methods are limited. Most require manual interventions or user-defined landmarks (3–6), leading to potential user biases and low throughput. Landmark-free automated approaches such as PASTE and GPSA (7,8) align slices mainly upon transcriptome profile similarities. However, gene expressions at a given location (aka, ‘spot’) are often obtained from a single or a few cells with low transcript detection rates (∼5%), yielding high noise in the expression readout. In addition, these transcriptome profiles can also vary due to non-spatial confounding factors such as cell state changes and batch variations, making it challenging to accurately determine spatial relationships across multiple slices using gene expression alone.

In this work, we approached this challenge by combining the gene expression with the microscopy data of the tissue slice that is acquired simultaneously with the transcriptome in many platforms (9). Our method, STaCker (Spatial Transcriptomics Common coordinate builder), formulates the CCF construction as an image registration task and provides an automated end-to-end solution without the need of predefined landmarks. Unlike previous approaches that relied on local image patches for spatial proximity assessment (8), whole-slide images were employed for both local and long-range information extraction to aid the alignment. Transcriptome data were converted into a contour map and overlaid onto the tissue image. The contour map emphasizes the spatial arrangement of cluster of tissue spots or cells, providing more resilience against data noise and batch effects.

We chose elastic models to register the derived composite image representations. Unlike rigid methods, elastic registration allows for local deformations such as stretches and contractions to better match the shape and structure of the objects in the images (10–12). This contrasts with rigid registration methods, which only allow for translations and rotations. Elastic registration is often used in medical imaging to align images acquired at different time points or from different modalities (11,13), and it has also been applied to the alignment of microscopic images (14,15). Recently, deep learning-based elastic registration methods, using convolutional neural networks (CNNs) to align images based on latent similarity embeddings, have gained popularity for their robustness and accuracy (16,17). We adopted the CNN framework in the development of STaCker.

Many deep learning algorithms learn how to properly perform tasks using acquired “training” data. Despite some success, this approach often suffers from a lack of sufficient training data; moreover, test cases containing features that are not present within the training data often cause model performance to suffer. A unique feature of STaCker is its exclusive use of synthetic data for training. It is especially beneficial for spatial transcriptome slice alignment, where available datasets are limited due to the nascent nature of the field and the high cost. Deep neural networks trained with synthetic images have been reported to be as effective or even more robust in some cases (18,19). Synthetic data allows control over data variability like color, contrast or rotation, aiding training a robust network. It can be easily augmented to improve model generalization and reduce overfitting. More importantly, a model trained with synthetic data is not tissue-specific, bolstering its overall applicability and generalization.

In the following sections, we present STaCker, a deep convolutional neural network based model that integrates the tissue imaging with gene expression data to learn feature representations and perform the alignment. STaCker is an automated end-to-end approach without the need of predefined landmarks. The performance of STaCker is assessed using simulated datasets with known ground truth as well as multiple real spatial transcriptome datasets, in comparison with multiple existing methods. STaCker effectively unified the spatial coordinates of deformed slices in various tissue types, surpassing existing methods. These outcomes demonstrate that STaCker is a valuable computational tool for establishing a CCF with spatial transcriptome data.

## Results

### Overview of the workflow of STaCker

STaCker is a deep learning algorithm designed to register a “moving” spatial transcriptome slice to a “reference” counterpart. The workflow involves several steps: processing tissue images, associating transcriptome information with each image, applying an image registration module to generate a deformation field, and using this field to align the moving spatial slice to the reference (Figure 1).

**Figure 1:**
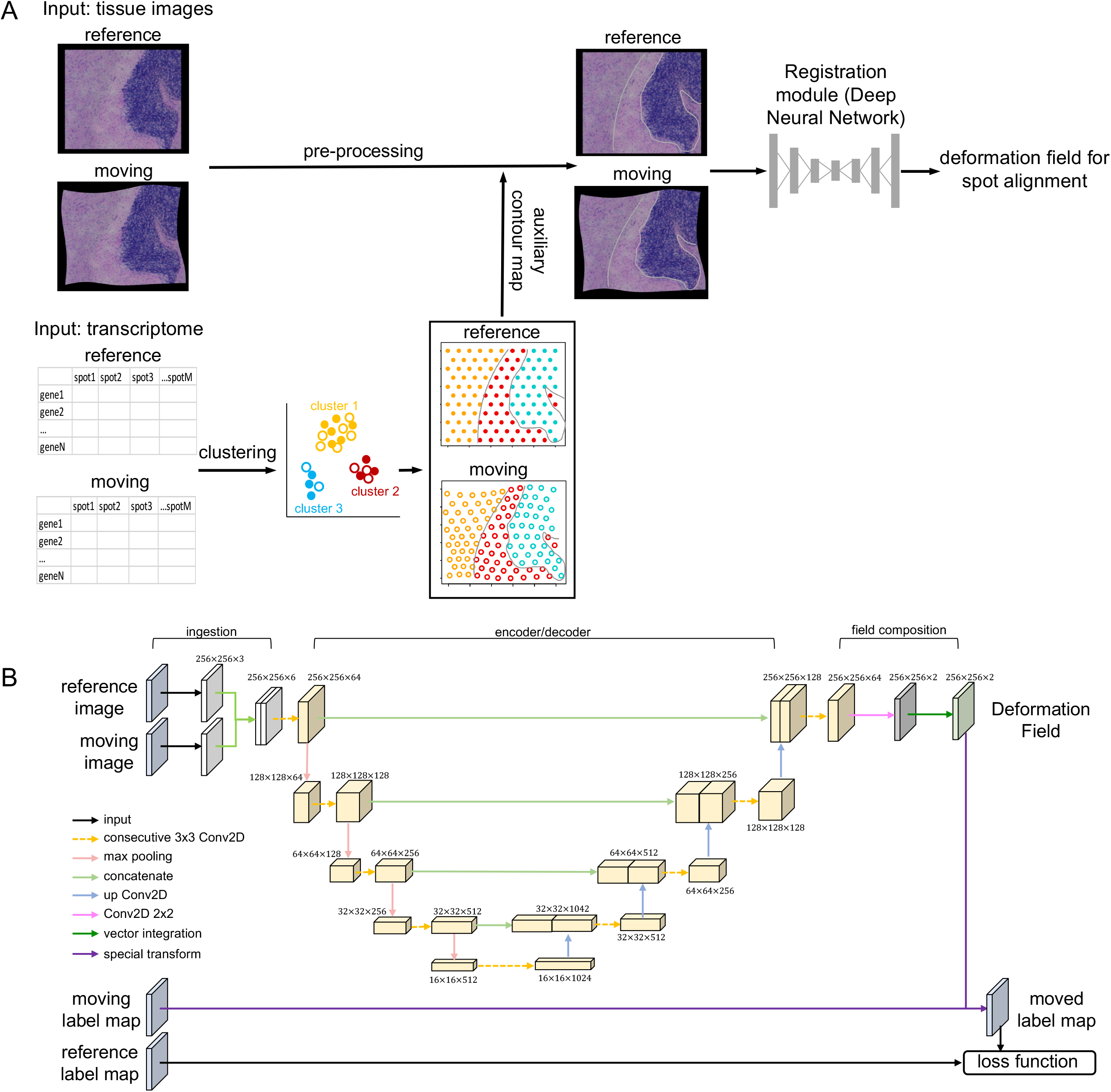
Schema representation of STaCker. The workflow (A) takes as inputs the tissue images from a pair of reference and moving spatial transcriptome slices, combined with the contour maps generated upon the gene expression profiles from the respective slices. The resulting composite images are subsequently aligned through a deep neural network based registration module. The registration module outputs the inferred deformation field to align the spatial coordinates of the spots/cells in the moving slice. The architecture of the registration module (B) takes a four-level contracting path and a four-level expanding path with skip connections at all levels. The final layer of the decoder is further convoluted to generate the spatial velocity field followed by a vector integration to output the deformation field for the alignment. Synthetic images with segmentation label maps (Methods) are used to train the module. The moved label map, after applying the deformation field to the moving label map, is compared to the reference label map. The difference constitutes the key component in the loss function.

The image data associated with each spatial transcriptome slice underwent a series of pre-processing step before being input into the alignment module (Figure 1A, top panel, see Methods for more details). These steps include color correction, non-tissue background masking, cropping, and resizing. Gene expression data from each slice was processed to create an image-like input (Figure 1A, bottom panel; Methods). Transcriptome profiles were normalized, transformed and integrated via a mutual nearest neighbor algorithm (20) to minimize batch biases. Dimension reduction and clustering identified groups of tissue spots or cells with similar gene expressions. The location of these clusters signified the spatial variation in molecular contents, providing valuable information about the tissue structure. A contour map derived from cluster boundaries was overlaid on the processed tissue image, which was input to the image registration module.

As shown in Figure 1B and detailed in Methods, the image registration module comprised an ingestion module, an encoder-decoder block, and a field composition module. The ingestion module employed a Siamese structure to receive the fixed reference image and the moving image. The encoder-decoder block implemented a U-Net backbone with skip connections at each level. The final layer of the decoder connected to the field composition module to produce the deformation field for alignment. While we primarily used the U-Net backbone in this study, it is essential to note that STaCker is not limited to this architecture. Other network models, such as visual transformers, capable of mapping features from imaging data to a deformation field, can also be utilized.

The deep neural network model was trained in a tissue-agnostic manner using synthetic data (Figure 2 and Methods). Each training instance consisted of four images: a colored reference image (*r̂*), a segmentation mask image (label map) associated with the reference image (*l*_*r*_), a colored moving image (*m̂*), and a label map associated with the moving image (*l*_*m*_). The label maps were created first, using Simplex noise distributions with customizable frequencies and amplitudes (Methods). Simplex noise is widely applied in generating natural appearing textures, including organ surfaces (21,22). It has also proven effective in synthesizing segmented microscopic medical images to augment training data for deep learning (23). Afterward, the colored reference and moving images were produced using the respective label maps as templates (Methods).

**Figure 2:**
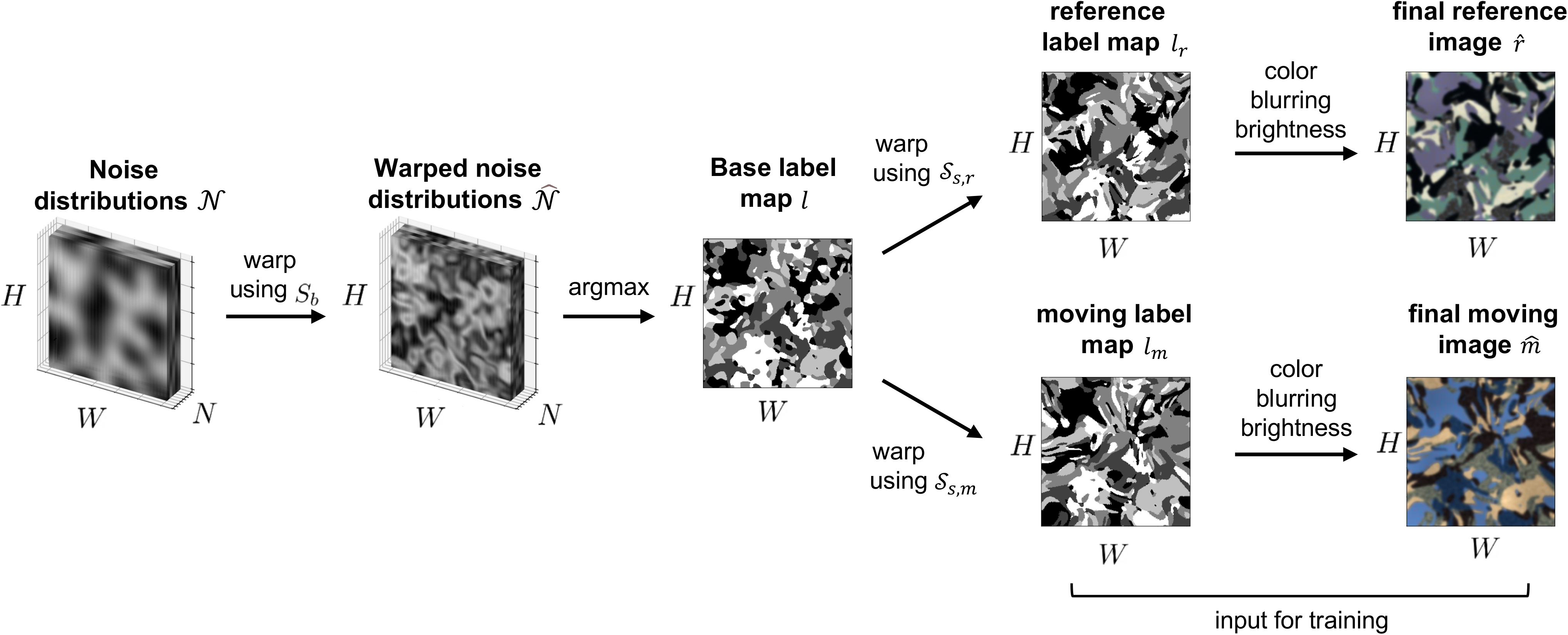
Generation of synthetic training data. The distribution *𝒩*contains *N* layers of 2-D Simplex noise of height *H* and width *W*. Each layer *n*_i_ corresponds to a class label *c*_i_. Sb denotes a collection of Simplex noise distribution *𝒮*_*b*,i_ of the same shape (*H*, *W*) that is applied to the layer *n*_i_ to form the warped noise distribution 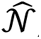 Next, 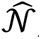 is condensed along the dimension *N* to form a 2-D base label map *l* of shape (*H*, *W*). Each pixel of *l* at the position (row, col) is assigned the class label *c*_i_ where layer *n*_i_ has the highest intensity at that position. *l* is further warped by separate Simplex noise *𝒮*_*s*,*r*_ and *𝒮*_*s*,*m*_ to create a reference label map *l*_*r*_ and a moving label map *l*_*m*_, respectively. Finally, the reference and moving RGB images, *r* and *m*, are generated based on the respective label maps *l*_*r*_ and *l*_*m*_ by assigning an RGB color to each of the class labels. The quadruplet *r*, *m*, *l*_*r*_ and *l*_*m*_ represents one sample in the training dataset.

The network model was trained using a loss function based on the Dice score between reference and moved label maps, along with a regularization term to discourage abrupt deformations (Eq. 1 and 2, Methods). The Dice score assesses the agreement of class labels instead of color and intensity, thereby accounting for the variations in fine-grained details like individual cell positions present in tissue slices. It emphasizes aligning cell regions rather than achieving precise cell-level alignment.

### Synthetic data training enables accurate image registration

We first assessed the performance of STaCker’s image registration module trained upon synthetic data, considering two tasks. First, we generated a new multi-color image not included in the training set and warped it either by adding Simplex noises of varying amplitudes (Figure 3A) or by random manual deformations. The manual warping was performed by the Warp Transform tool in GIMP (24) with a brush size of 25% of the largest dimension of the image to drag around arbitrary regions in random directions. The manual distortions represented independent warps that the module did not encounter during training. The distortion levels were quantified using normalized cross correlation (NCC, Eq. 3), a metric that measures the similarity between the warped and the unwrapped images. For comparison, we included the results from the affine alignment and the nonlinear transformation offered by a state-of-the-art image registration tool ANTs (25). After the image registration, STaCker significantly improved the NCC scores (p-values <0.001, Figure 3A) in all warped cases, consistently outperforming ANTs and the affine alignment by a large margin.

**Figure 3:**
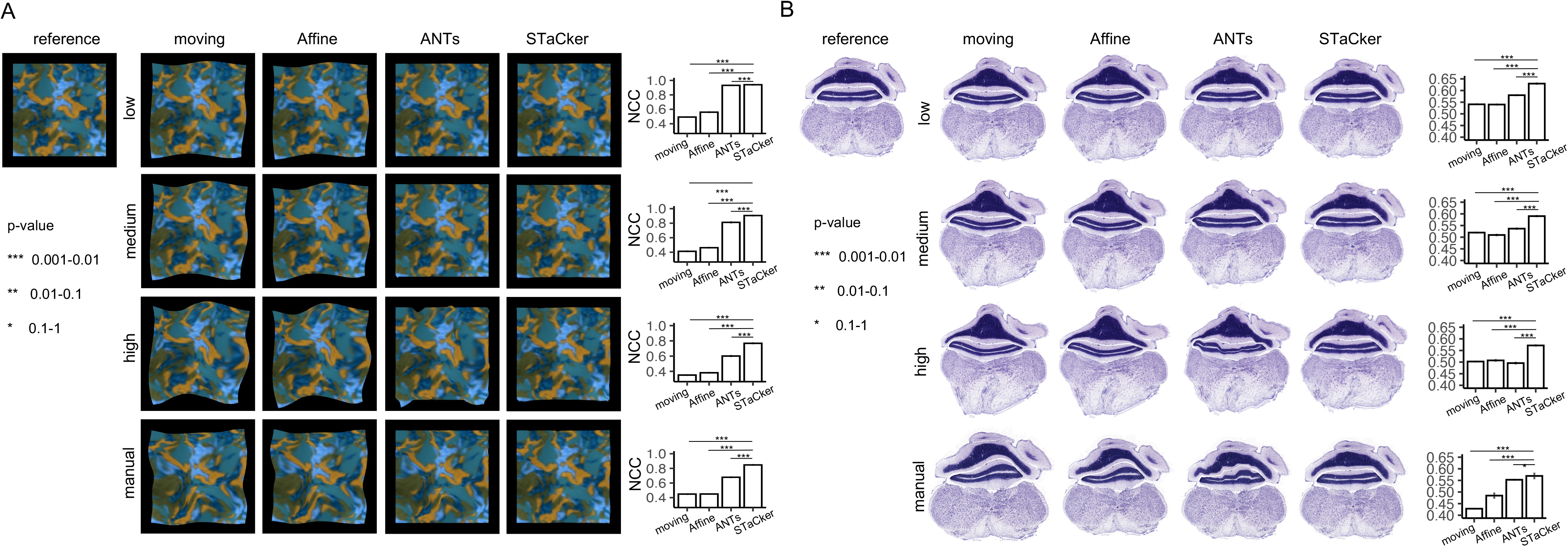
Validation of the image registration module of STaCker trained by synthetic data. A synthetic image not used for the training (A) or a real tissue histology image (B) was digitally distorted to a low, medium or high level, or manually warped to generate a series of moving images. The original unwarped images served as the references. The degree of the distortions is quantified as the “moving” NCC score in the bar plots. The image module of STaCker, together with the Affine method and the non-linear alignment (“SyN”) method offered by ANTs, were applied to align each of the moving images to the reference. The aligned images from each method are displayed, together with their post-alignment NCC scores shown in the bar plots. The displayed NCC value of STaCker and ANTs is the mean over 10 repeated runs. The associated error bars are also plotted yet are very small. The statistical significance of the increase in STaCker’s NCC values relative to other methods is marked by asterisks and explained in the annotation.

In the second task, we considered an actual tissue H&E staining image. A coronal slice of mouse hindbrain obtained from Allen Brain Institute (see Data availability) was used as the reference image. We applied Simplex noise wrapping and manual wrapping to create a set of moving images (Figure 3B). Like the results in Figure 3A, STaCker outcompeted ANTs in all test cases. Note that ANTs sometimes introduced wavy edges around the pyramus and medulla regions. In contrast, these apparent artifacts were absent in STaCker. The result demonstrates that STaCker can effectively register real tissue images, despite training solely on synthetic images.

### STaCker accurately aligns digitally warped spatial transcriptome slices

Upon the validation of the image registration module, we proceeded to evaluate STaCker’s ability to align spatial transcriptome slices. We chose a sagittal dissection of mouse brain profiled using Visium (see Data Availability) and digitally warped it to various degrees (Figure 4). The level of distortion is quantified using the NCC and Mean Square Error (MSE, Eq. 4) scores associated with the applied Simplex noise warping. The undistorted slice and each distorted slice formed a pair of reference and moving inputs for the model, respectively. The advantage of digital warping is that the ground truth is known. A total of 3248 in-tissue spots that passed quality control were included.

**Figure 4:**
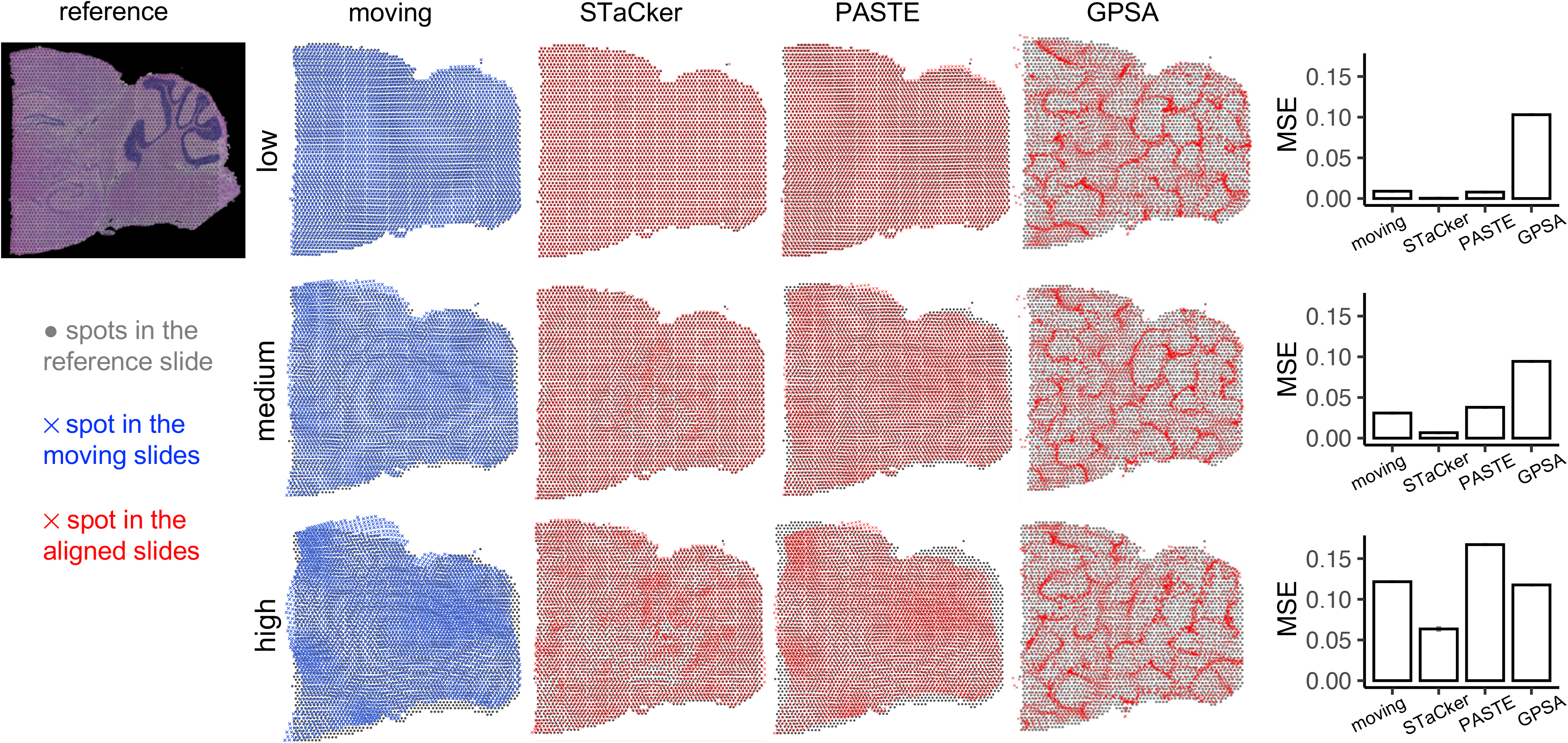
Evaluation of STaCker in aligning digitally warped spatial transcriptome slices of mouse brain. The reference is a mouse sagittal posterior brain slice profiled by 10x Genomics Visium platform. It was digitally warped using Simplex noises to a low (noise amplitude = 5, NCC of the deformed image=0.606), medium (noise amplitude = 10, NCC of the deformed image=0.566), or high (noise amplitude = 20, NCC of the deformed image=0.534) level to generate a series of moving slices (noise frequency remains 1 for all warping). The discordance between the spatial coordinates of the spots in each moving slice and those in the reference slice is quantified by the MSE (Methods) shown in the bar plots. STaCker, together with previously published methods PASTE and GPSA, were applied to align each of the moving spatial transcriptome slices to the reference. The coordinates of the spots either before (blue crosses) or after the alignment (red crosses) are displayed together with the reference spot coordinates (gray dots) to aid the visual comparison. The post-alignment MSEs from each method are illustrated in the bar plots. Values from STaCker and GPSA are the average over 10 runs, shown with small error bars.

Figure 4 summarizes the alignment of the digitally warped moving slices to the reference slice performed by STaCker as well as PASTE and GPSA (7,8), as all three are landmark free alignment programs. As the warping distortions increased, the MSEs between the pre-alignment moving slices and the reference slices also increased (0.0089, 0.0308 and 0.1216, Figure 4). In each case, STaCker significantly reduced the MSE (0.0003, 0.0067 and 0.0635, respectively), accompanying the good matching between the post-alignment (red crosses) and the reference coordinates (gray dots). The residual discordance in the post-alignment of STaCker increased slightly with larger initial distortions. This phenomenon is expected as STaCker employs a regularized loss function. Nevertheless, even in the most discordant case (Figure 4, bottom row), STaCker placed the majority (73.1% on average) of the spots associated with the highest MSEs (top 10 %) nearest to a spot in the reference slice that has the matching class label, indicating these spots were correctly located in the expected regions for their class labels (Supplementary Figure 1). Thus, although STaCker might not perfectly align these most dislocated spots, it managed to assign them biologically correct locations.

Both PASTE and GPSA were executed using the author-exemplified parameter settings for Visium slice. PASTE performed less well with substantially higher post-alignment MSEs in all cases. GPSA also yielded a higher MSE than STaCker, though lower than PASTE. Additionally, the aligned tissue spots by GPAS appear to aggregate upon convergence (see also Supplementary Figure 2A). This result persisted when running GPSA with different choices of parameters (Supplementary Figure 2C). We speculated that GPSA may suffer a low sensitivity to transcriptomically differentiate the neighbor spots given the limited number of genes (10 by default) it handles. Increasing the number of genes to 100 did not resolve the aggregation (Supplementary Figure 2B-C). Attempts of more genes led to a memory shortage issue that rendered GPSA computationally impractical and thus were not pursued further.

STaCker is also suitable for use with in situ hybridization (ISH)-based spatial transcriptome data, such as MERFISH and Xenium (26,27), where images of cell body or nucleus staining, like DAPI (4ʹ,6-diamidino-2-phenylindole) labeling, are often available from the tissue. We evaluated STaCker against PASTE and GPSA using a coronal mouse brain slice profiled by the 10X Genomics Xenium platform (see Data availability). Our observations indicated that STaCker significantly reduced the pre-alignment MSE while other methods failed to do so (Supplementary Figure 3). This suggests that STaCker maintains superior performance in ISH-based profiling data. Unlike sequencing-based spatial techniques used above (e.g. 10x Genomics Visium) that profile the whole transcriptome, ISH-based platforms typically profile only a few hundred genes, making it more challenging to differentiate a cell from similar ones based solely on these gene expressions. In this test, both PASTE and GPSA were unable to reduce the MSE post-alignment. These findings underscore the benefits of properly utilizing the image modality, as demonstrated by STaCker, in both sequencing- and ISH-based spatial profiling, which encompasses most spatial transcriptome applications.

### STaCker’s leading performance is tissue agnostic

As STaCker is trained on synthetic data, it is expected to be applicable to images of various tissue types. We conducted a benchmarking alignment in human lymph nodes (see Data Availability) in a similar manner to the previous evaluation. The tissue texture in the human lymph nodes is noticeably different from the mouse brain (Figure 4 and Figure 5, leftmost columns).

**Figure 5:**
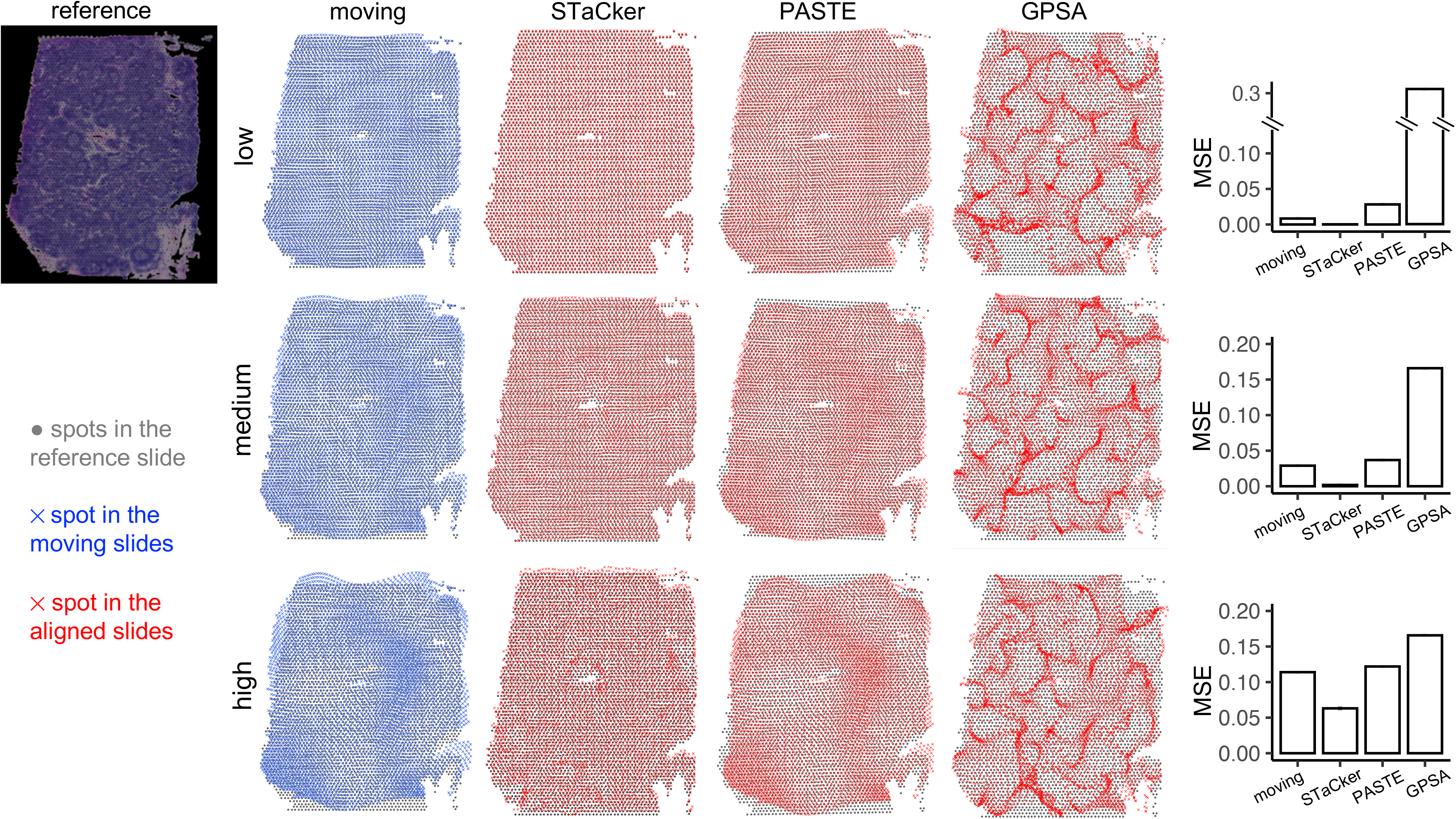
Performance of STaCker in aligning digitally warped spatial transcriptome slices of human lymphnodes. The reference is a human lymphnode dissection profiled by 10x Genomics Visium platform. It was digitally warped using Simplex noises to a low (noise amplitude = 5, NCC of the deformed image=0.621), medium (noise amplitude = 10, NCC of the deformed image=0.540), or high (noise amplitude = 20, NCC of the deformed image=0.504) level to generate a series of moving slices (noise frequency remains 1 for all warping). The discordance between the spatial coordinates of the spots in each moving slice and those in the reference slice is quantified by the MSE (Methods) shown in the bar plots. STaCker, together with previously published methods PASTE and GPSA, were applied to align each of the moving spatial transcriptome slices to the reference. The coordinates of the spots either before (blue crosses) or after the alignment (red crosses) are displayed together with the reference spot coordinates (gray dots) to aid the visual comparison. The post-alignment MSEs from each method are illustrated in the bar plots. Values from STaCker and GPSA are the average over 10 runs, shown with small error bars.

As shown in Figure 5, there are three moving slices that exhibit increasing distortions compared to the reference slice. STaCker, PASTE and GPSA were applied to align each the moving slices to the reference slice, following the same procedure as described earlier and in Methods. After the alignment, STaCker achieved a substantial reduction in MSE in all moving slices (rightmost columns in Figure 5, from 0.0084 to 0.0002, 0.0289 to 0.0017, 0.1139 to 0.063, respectively). Compared to PASTE and GPSA, STaCker delivered the lowest MSEs and the highest consistency between the post-alignment coordinates and the reference coordinates (Figure 5, grey dots and red crosses). The result suggests that STaCker retains its leading performance regardless of the tissue type.

### STaCker outperforms in the de novo alignment of slices

STaCker allows users to align multiple slices from repeated experiments by defining one of the slices as the reference for aligning the remaining, known as the “template-based” alignment. Alternatively, STaCker offers a “template-less” option where slices are aligned to each other without a single fixed reference, also referred to as de novo alignment in this work. To evaluate STaCker’s performance in de novo alignment, we used a section from the previously discussed mouse sagittal posterior brain slice and digitally warped it four times independently to create a set of slices for alignment. The section contains 115 tissue spots. STaCker was run in the “template-less” mode (see also Methods), PASTE in its “center-alignment” mode, and the four slices were passed to GPSA as four “views”.

Before alignment, the average MSE over every pair of slices is 0.110 (Figure 6, the panel of unaligned coordinates). After de novo alignment, STaCker reduced the mean pair-wise MSE to 0.046, followed by 0.105 by PASTE and 0.55 by GPSA. The four sets of post-alignment coordinates of the spots by STaCker were most mutually consistent as well (Figure 6, the panel of STaCker aligned coordinates). Note that in STaCker’s template-less mode, none of the slices retained its original spot coordinates. Instead, they converged towards each other as expected in de novo alignment.

**Figure 6:**
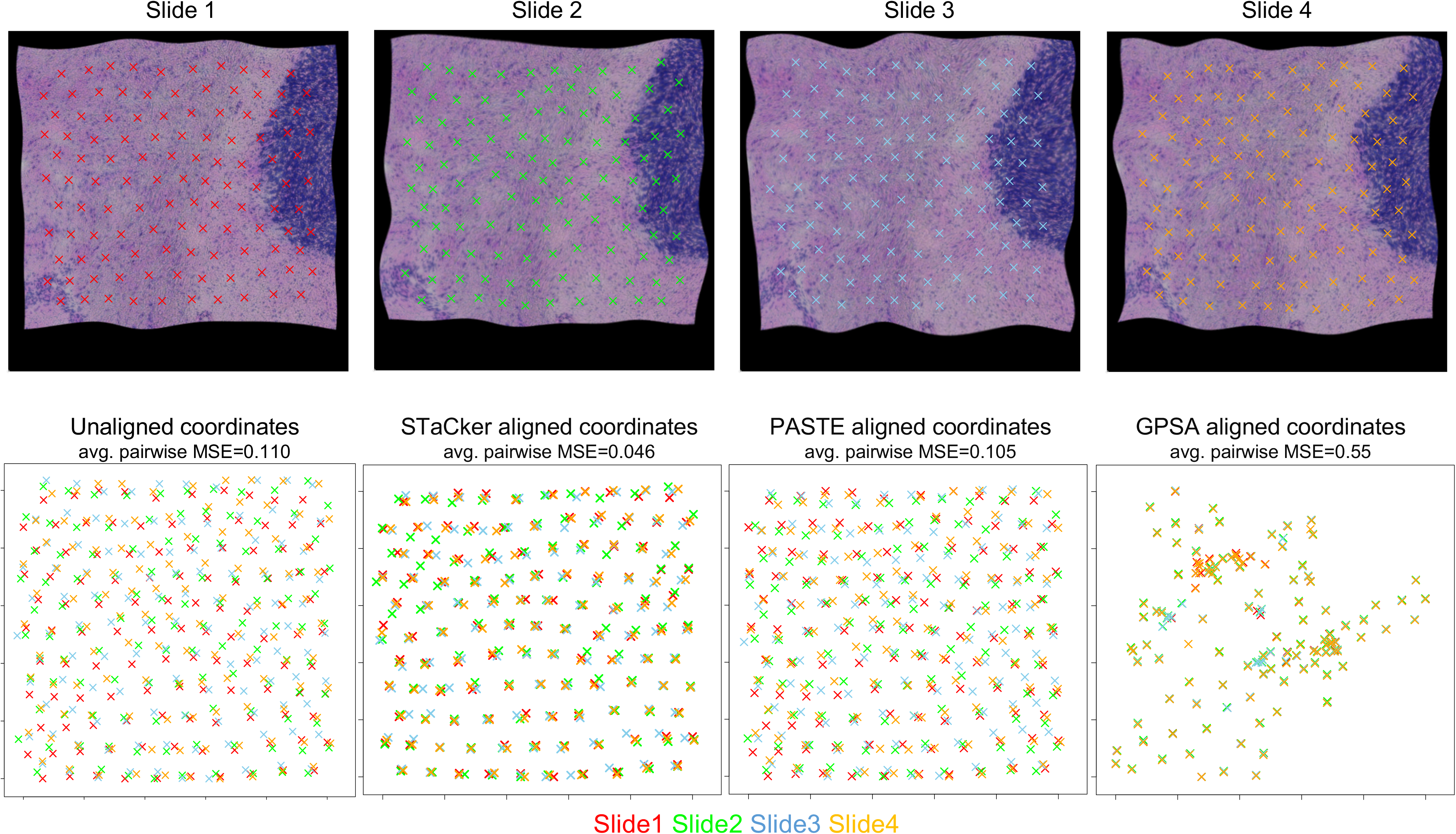
Performance of STaCker in the de novo alignment of spatial transcriptome slices. The top row displays four moving spatial transcriptome slices that were independently warped from a reference slice taken from the mouse posterior brain used in Figure 4, with the spot coordinates shown as crosses over the tissue images (slice 1: red, slice 2: green, slice 3: blue, slice 4: orange). The warping was conducted using random-seeded Simplex noises with an amplitude of 15 and frequency of 1. The mean pair-wise NCCs among the tissue images of the moving slices is 0.198. The average pair-wise MSE among the spot coordinates in the moving slices is 0.10. The bottom row illustrates the spot coordinates from four slices before the alignment (“Unaligned coordinates”) and after the alignment by STaCker, PASTE and GPSA, respectively, using the same colors and cross symbols as shown in the top row. The post-alignment average MSE over all pairs of slices is 0.046, 0.105 and 0.55 for STaCker, PASTE and GPSA, respectively.

The four slices aligned by PASTE remained considerably discordant. The spots aligned by GPSA aggregated and largely dislocated from their original positions (Figure 6, the panel of GPSA aligned coordinates). On average 0.4% of the spots in a slice retained the same neighboring spots within 150 microns (1.5 fold of inter-spot distance in these slices) after the alignment, indicating GPSA did not maintain the integrity of the spatial relationships among the spots. In contrast, both STaCker and PASTE preserved the spatial neighborhood well (over 97% of spots retained their neighbors). The results of the three programs echo their performance in aligning a single moving slice to a reference, which is expected as the same underlying algorithms are used for either the pair-wise or the de novo alignment in each program.

### Application of STaCker in real-world data

STaCker’s robust performance in the previous benchmarking encouraged us to apply it to spatial transcriptome slices with real deformations. We considered two most common scenarios: slices acquired from adjacent tissue dissections (Figure 7A), or from several biological replicates (Figure 7B). Figure 7A displays two consecutive sagittal slices of a mouse brain profiled using Visium (see Data Availability). They have a mild but noticeable discoordination. For instance, slice 2 appeared to be horizontally more stretched and slightly shifted upwards compared to slice 1 (Figure 7A, unaligned tissue images).

**Figure 7:**
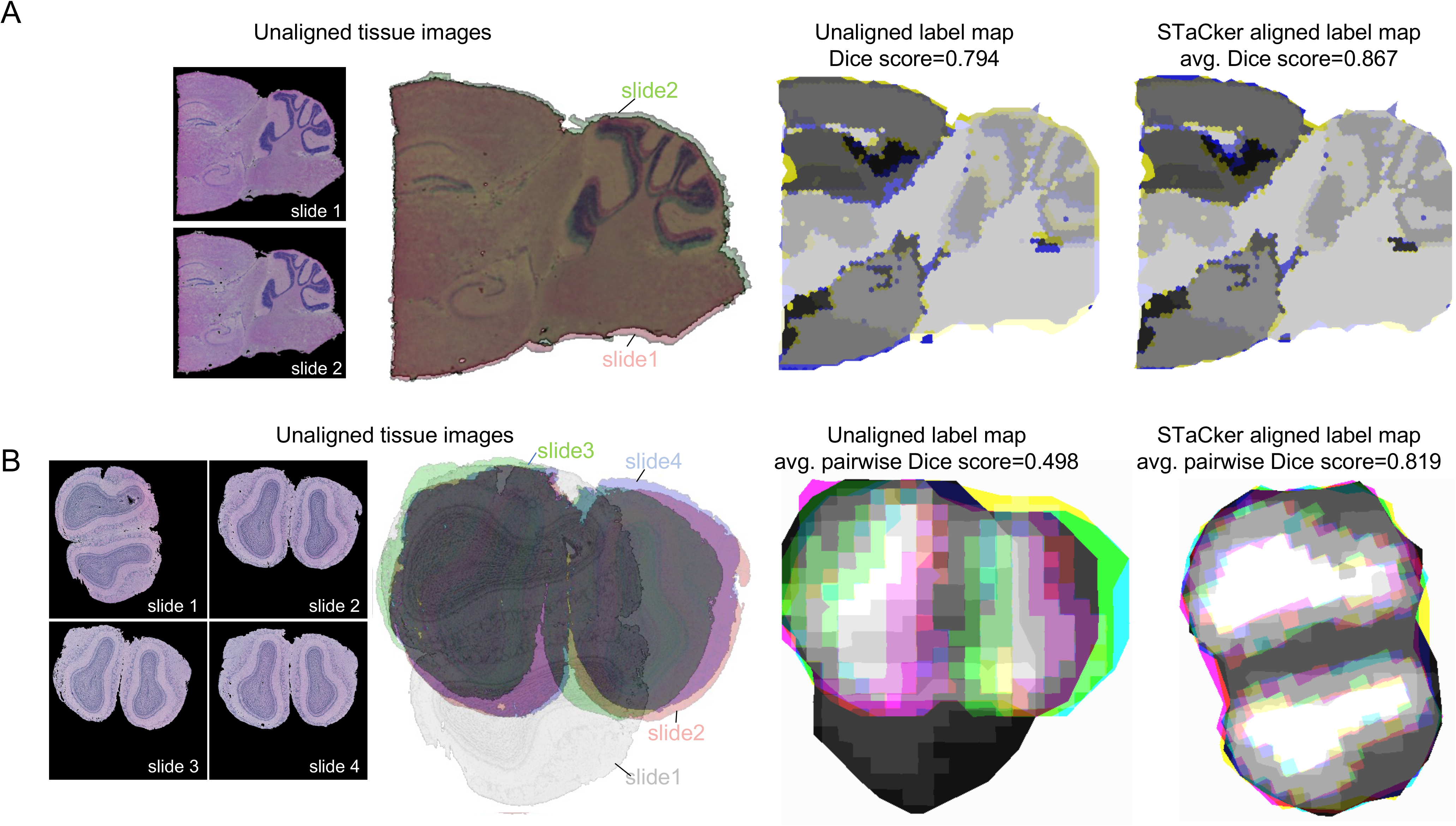
Application of STaCker in real spatial transcriptome slices. A): The first panel shows two consecutive sagittal dissections of mouse brain profiled by Visium. Prior to the alignment, the difference between the slices is illustrated in the superimposed tissue images (the second panel, red for slice 1 and green for slice 2). In addition, to assess the spatial consistency of tissue spot clusters between the slices before (the third panel) and after (the fourth panel) the alignment, we present them in the form of superimposed class label maps. The slice 1 and slice 2 contain identical classes, depicted in distinct shades of yellow and blue, respectively. The superimposed map is generated by combining the colors from both slices in each RGB channel with a mixing ratio of 0.5 each. In this map, regions displaying various shades of grey indicate agreement between the class labels of the two slices. The accompanying Dice score between the unaligned label maps is 0.794. After the alignment of STaCker, the Dice score increases to 0.867. B): The first panel displays four biological replicates of dissected mouse olfactory bulbs. To illustrate their spatial discordance, the second panel showcases superimposed tissue images with colors representing each slice (grey for slice 1, red for slice 2, green for slice 3, and blue for slice 4). The classes in the label map of the slice 1, 2, 3 and 4 are represented in varying shades of grey, red, green and blue, respectively. For the superimposition of label maps, the colors from all four slices are combined in each RGB channel with an equal mixing ratio of 0.25. Regions exhibiting different shades of grey signify agreement between the class labels across the four slices. Prior to STaCker’s alignment, the Dice score is 0.498 (shown in the third panel). After STaCker’s alignment, the Dice score improves to 0.819 (shown in the fourth panel).

As the ground truth alignment of the spot coordinates was unknown, we evaluated the spatial concordance of the clusters of spots between the slices. The tissue spots from both slices were classified into 12 clusters based on their transcriptome profile. Each pixel at the centroid of a spot received a class label corresponding to the cluster of that spot. Other pixels were assigned a class label according to their nearest spot centroid, resulting in a label map. To improve visualization, the label map of a slice was rendered using various shades of a single color, with each shade representing a different class. The color selection was designed to ensure that when label maps were overlaid, only areas with matching class labels across the slices appeared in shades of achromatic gray, while non-matching regions retained their distinct colors from the respective slices. As shown in the superimposition of the unaligned label maps in Figure 7A, there were several areas with mismatching class labels before the alignment, typically around the edges of tissue substructures. For instance, numerous blue or yellow shades were noticeable along the cerebellum granular layers. The inconsistencies were greatly reduced after the alignment by STaCker, as evidenced by the reduction of the blue or yellow regions and the corresponding increase of grey shades (Figure 7A, STaCker aligned label map). The improvement in the class label maps was quantified by a notably elevated mean Dice score from 0.79 to 0.87.

The second dataset contained four biological replicates of the mouse olfactory bulb slices published earlier (1). There were slight to large discrepancies in the slice orientation and shape with occasional tears (Figure 7B, unaligned tissue images), as well as non-negligible batch effects in the transcriptome profiles (Supplementary Figure 4A). In Figure 7B the unaligned label map illustrates the extent of the spatial consistency among the original slices. The label maps were generated using the same method outlined in Figure 7A. Pixels that had a consistent class label across all slices were depicted in different shades of gray as well as white. On average, these pixels only made up 6.2% of all in-tissue pixels on each slice. The corresponding pair-wise Dice scores that describe the coordinate consistency among the clusters of spots averaged 0.498. We applied STaCker to these slices in the template-less mode. The pair-wise Dice scores significantly increased to 0.819 (student t-test p-value= 0.0093) in conjunction with the improved percentage of matching areas across all slices (57.5% on average), indicating an effective harmonization of the slices.

## Discussion

Constructing a unified coordinate system across multiple spatial transcriptome slices is an important but unmet need for data comparison, integration, and interpretation. A common approach involves adjusting the spatial proximity of tissue spots or cells based on the similarity of their gene expression profiles, within a permitted range of translocation. Nevertheless, this can be challenging due to the noisy nature of gene expression at the single cell or near single cell resolution, and the potential for significant expression alterations due to biological or technical batch effects.

STaCker addresses these challenges by treating the construction of a common coordinate as an image registration problem. Unlike previous efforts that used the image patches associated with each spot to identify the spot-to-spot similarity and their spatial closeness (8), STaCker convolutes the image features over the entire slice, capturing better contextual information and global features to aid alignment. STaCker also converts the transcriptome data into a contour map and superimposes it with the tissue image. The contour map captures the spatial organization of the clusters of spots/cells defined based on the dimensionally reduced gene expressions, making it less susceptible to data noise and batch effect. The molecular level transcriptomic readout provides supplementary information to the tissue image, especially in areas where the image may not be sensitive to changes in the tissue microenvironment due to limited resolution or unperturbed morphology.

STaCker’s image registration module was trained using synthetic images of arbitrary colors and shapes, aiming to build a general model that is agnostic to specific tissue morphologies. This approach has proven successful given that STaCker worked well with various tissue types. The use of synthetic images also addresses the challenge of limited training data when building a deep neural network for image registration. During the training, STaCker tailored its objective to the application of spatial transcriptome slices. The loss function prioritized the consistency between the images regarding the segmentation labels rather than their pixel-wise details. This is because we do not anticipate profiling identical cells or the same composition of them across slices. Consequently, STaCker may ignore aligning some subtle patterns in an area if the region is deemed homogeneous given the tissue image and the gene expressions. We believe the output in such scenarios remains biologically meaningful because the cells/spots within these areas are considered indistinguishable upon the available data, implying no further alignment is necessary. If subregions can be identified with higher resolution images and/or more sensitive transcriptome data, STaCker can adapt to these enhancements without altering its framework.

In the current work, the primary application of STaCker is to unify the spatial transcriptome coordinates among multiple replicates or several consecutively dissected slices. It can be applied in two modes, either aligning to a user-defined reference or mutually among the inquired slices themselves. Our benchmarking showed that STaCker can effectively improve the deformations. Nevertheless, the larger the distortions the less extent STaCker can restore given applied regularizations. On the other hand, its conservative behavior may be desired in practice to avoid over correction.

Our results demonstrate the effectiveness of the image-guided strategy in constructing a common coordinate framework for spatial transcriptome slices. This work also provides a useful foundation for future improvements. For example, we aim to enable STaCker to handle full megapixel resolution images acquired in spatial transcriptome measurements that preserve more subtle changes in the tissue microenvironment. Multi-scale training and workflows to enhance parallelization and scalability may be considered. In addition, it is critical to differentiate slice distortions from true structural changes, such that the spatial profiling across different biology states can be unified and compared. We expect a better digestion of the rich morphology features, in conjunction with fine-tuned regularizations, will help address these challenges and further broaden the application of STaCker.

## Methods

### Input image data pre-processing

The image associated with a spatial transcriptome slice was preprocessed using Python package histomicstk (28). For histology staining images, we applied the Reinhard color normalization algorithm if noticeable color deviations or variations existed. The non-tissue background of each image was masked using saliency.tissue_detection module. If it was challenging to identify non-tissue background pixels using the colored image, an intensity threshold was applied to the grayscale version of the image, followed by the restoration of the color in the identified tissue foreground. Each color channel was subsequently scaled to be within [0,1]. The image was cropped to remove undesired background sections and resized to a given resolution for input into the model.

### Input transcriptome data pre-processing

The gene expression data from a spatial transcriptome slice was converted into an image-like input as the following (Figure 1A, bottom panel). For each slice, the transcriptome profiles of all spots (or cells) were collected into a count matrix. The columns of the count matrix represented the spot or cell IDs and the rows represented genes. The count matrix was normalized to have an equivalent total number of counts per column, followed by natural log transformation and standardization using R package Seurat (20). After processing each slice, the transcriptome data from all slices were integrated to minimize batch biases using Seurat, followed by a dimensionality reduction. The top 30 principal components (PCs) were used to perform a mutual nearest neighbor-based clustering to identify major clusters of the tissue spots or cells.

After identifying the clusters, the pixel at the center of each spot or cell was was given the corresponding cluster label of that particular spot or cell. The labels for the remaining pixels were estimated using a Voronoi partition created from these central pixels. We then extracted the boundaries between different cluster regions, creating a contour map that represented the spatial organization of the tissue spots or cells based on the transcriptome. Pixels on the contour map were given an intensity of 1 along the boundaries in each color channel. This contour map was then overlaid on the previously processed image, specifically at the pixels underlying the contour lines, with a blending ratio of 0.3:0.7. These composite images were then fed into the image registration module. Users have the flexibility to bypass the inclusion of the transcriptome-derived contour map when its utility is considered limited, such as when the transcriptome data quality is poor, or when the transcriptome provides little or no extra information about the spatial organization of cells.

### Deep Neural network architecture

As depicted in Figure 1B, the registration model was composed of an ingestion module, an encoder-decoder block and a field composition module. The ingestion module uses a Siamese structure to receive the fixed reference image and the moving image to be aligned.

The encoder-decoder block is embodied in a U-Net backbone (29), consisting of four levels of contractions and four levels of expansions with skip connections included at every level. The final layer of the decoder is connected to the field composition module to output the deformation field for the alignment as well as its inverse for downstream convenience. Note that the model is not limited to a U-Net architecture. Other network models that can extract and map features from imaging data to a deformation field can be used.

### Model training using synthetic images

The model was trained in a tissue-agnostic manner using synthetic training data. In that way, the alignment model can apply to any tissue type and/or image acquisition protocols (e.g. various histology or fluorescence microscopy staining techniques). Additionally, using synthetic training data can help address the issue of limited training data availability which is common in deep neural network training. In contrast to existing technologies, our approach to image synthesis and training is uniquely devised to suit the task of aligning spatial transcriptome slices (Figure 2). Each instance of a training data record contains a quartet of synthetic images: a colored reference image (*r̂*), a segmentation mask image (referred to as the label map in this work) associated with the reference image (*l*_*r*_), a colored moving image to be aligned with the reference image (*m̂*), and a label map associated with the moving image (*l*_*m*_). Pixels in the label map are classified into a finite number of distinct classes, with each pixel being assigned a specific class label. Although those classes are abstract regarding the training data, they can reflect spatially segregated regions of tissue microenvironment and/or cell compositions in the context of a tissue slice.

More specifically, for a particular quartet of synthetic images of dimension (*H*, *W*), where *H* denotes the image height and *W* denotes the image width, the reference and moving label maps were first generated. This process began with generating a multi-layer noise distribution *𝒩* of shape (*N*, *H*, *W*), where *N* denotes the number of layers and is customizable. Each layer *n*_i_ | *i* ∈ {0,1, …, *N* − 1} corresponded to a class label *c*_i_. Each layer *n*_i_ was a single channel image of smooth textures created using random seeded two-dimensional Simplex noise (OpenSimplex version 0.4.2)(30) with the maximum amplitude of 0.5 and a frequency uniformly sampled from [5/512, 15/512]. Every layer *n*_i_ of the distribution *𝒩* was then individually warped by a Simplex noise *𝒮*_*b*,i_ of the same shape (*H*, *W*). When generating the warping noise, we applied random seeding with a frequency between 15/512 and 25/512 and a maximum amplitude sampled log-uniformly from 10.24 to 102.4. The translocations defined by *𝒮*_*b*,i_ indicate for each pixel of the warped layer *i* which pixel intensity from the original layer *i* to take. This step formed the warped noise distribution 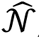 also of shape (*N*, *H*, *W*).Next,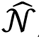 was condensed along the dimension *N* to form a “base” label map *l* of shape (*H*, *W*) whose elements consist of the class labels *c*_&_, *c*_1_, …, *c*_*N*)1_. In other words, any arbitrary element of *l* at a position (*row*, *col*) was defined by finding the solution to *l*(*row*, *col*) ≡ *c*_i_ = arg max 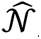 (*i*, *row*, *col*) | *i* ∈ {0,1, …, *N* − 1} (Figure 2). An additional copy of *l* was made, and i each copy was warped further by an additional separate Simplex noise *𝒮*_*s*,*r*_ and *𝒮*_*s*,*m*_ parameterized with a frequency sampled from [95/512, 105/512] and a maximum amplitude sampled from [10.24, 102.4]. This formed a “reference” label map *l*_*r*_ and a “moving” label map *l*_*m*_, respectively, each of which looks similar in appearance but exhibits noticeable differences (Figure 2).

We then converted the label maps *l*_*r*_and *l*_*m*_[shape (*H*, *W*)] into reference and moving images *r* and *m* [shape (*H*, *W*, 3)], respectively. To do this, we randomly selected an RGB color for each class *c*_i_ that appeared within the label maps and assigned that color to all pixels belonging to class *c*_i_ in images *r* and *m*. Both images were then subjected to Gaussian blurring and an intensity bias field to introduce variations in image sharpness and illumination that might appear in realistic microscopic images. This culminated in colored reference and moving images *r* and *m*, respectively.

The network model was trained concerning a loss function based on a similarity metric of a pair of reference and moving label maps and a deformation vector field to align the moving image to the reference image (Figure 1B):

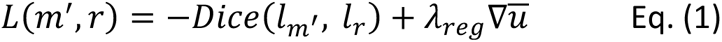

where *r*, *l*_*r*_ represent a training reference image and the reference label map associated with that image, respectively. *m*^+^, and *l*_*m*_′ represent, respectively, a moved image and the moved label map associated with that image, after applying the inferred deformation field by the model. *λ*_*reg*_ is a regularization factor, and 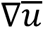 is the gradient of the inferred deformation vector field. In Eq. (1), the regularization term 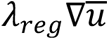 discourages abrupt large deformations. The Dice score *Dice*(*l*_*m*_′, *l*_*r*_) is a similarity metric that assesses agreement of the pixel-wise class labels between the reference label map and moved label map, instead of the pixel-wise color and intensity agreement between the reference image and the moved image.

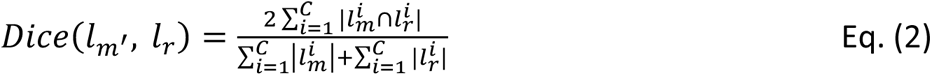

where *C* is the number of classes, 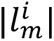 and 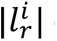 denote the number of pixels assigned class *i* in *m r l*_*m*_ and *l*_*r*_, respectively. 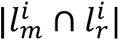 represents the number of pixels that are simultaneously assigned class *i* in both maps. This design of the loss function suits the fact that the tissue slices to be aligned are not expected to be identical regarding fine grain details, such as the position of individual cells. Instead, the preferred matching is regarding the regions of cells. The training was conducted using mini-batches of 32 images and the Adam optimizer. The learning rate began at 1e^-4^ and was decreased exponentially whenever the loss reached a plateau. Convergence was achieved within 300 epochs.

### Alignment of spatial slices

The trained STaCker model can align the coordinates of spots/cells in a moving slice to a reference slice. It takes as input the tissue images or the respective transcriptome-incorporated images and outputs a deformation field. The deformation field is resampled at the coordinates of the tissue spots or cells on the moving slice before it is used to align the spots with the reference slice.

STaCker uses a tessellation strategy to manage input images that exceed the neural network’s default input size (256×256 pixels). It generates a uniform tiling along the width and the height of the reference (or the moving) image, with each tile image fitting the default size of 256×256 pixels. Each tile partially (default 25%) overlaps spatially with every other adjacent tile image. Each moving tile image matches one and only one reference tile image. The matching pairs are supplied to the registration model and yield *N_T_* tile deformation fields, where *N_T_* denotes the total number of tiles in the reference image which is equivalent to that of the moving image. Each of the *N_T_* tile deformation fields and every one of its adjacent tile deformation fields share a defined common region. STaCker joins the *N_T_* tile deformation fields to form a full-size deformation field by calculating a weighted average of the tile deformation fields overlapping at each pixel within the common region. The weight is inversely proportional to the distance between the center of a tile and the specific pixel.

Users have the flexibility to run STaCker multiple times if the moving slice has a significant displacement. This can be done until the spatially transformed coordinates of the spots/cells stabilize or a user-specified number of iterations is achieved (e.g., 3 iterations). All the STaCker results presented in this study were obtained from a single iteration, which yielded favorable results compared to other tested methods (see Results). While additional iterations may not generally be required, we provide this option for added flexibility.

STaCker can align a group of slices either in a “template-based” mode or a “template-less” mode. In the template-based mode, one of the slide slices is set as the fixed reference, and the remaining slide slices are aligned to it, using the same process of aligning a pair of slides as described earlier. On the other hand, the “template-less” mode first adjusts the scale, position, and orientation of the slices in a coarse-grained manner, wherein one user-specified slice is taken as the reference, and the rest are affine-aligned to it. Subsequently, for each slice *s*_i_ among the *S* total slices, STaCker performs a pairwise alignment by applying the trained registration model on *s*_i_ using each of the (*S* −1) remaining slices *s*_j_(*j* ≠ *i*) as fixed references. The resulting (*S* −1) sets of aligned coordinates of *s*_i_, together with the unaligned coordinates of *s*_i_ (which can be considered as the result from the alignment against itself), are averaged and output as the final post-alignment coordinates of the slice *s*_i_.

### Quantitative evaluation of the alignment

The normalized cross correlation (NCC) is a measure of the consistency of pixel-wise intensity distributions between a pair of images. The NCC between two images R and M, is calculated as follows:

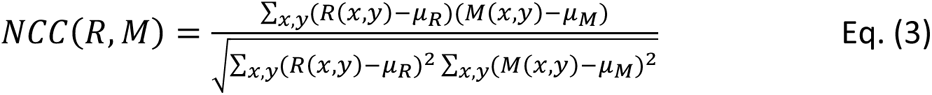

Here *R*(*x*, *y*) and *M*(*x*, *y*) are the pixel intensities of images *R* and *M* at coordinates (*x*, *y*), respectively. *μ*_*R*_ and *μ*_*M*_ are the mean intensities of images *R* and *M*. Note the formula is simplified to suit our application, where both images consistently possess identical dimensions and expect full overlap.

Mean Squared Error (MSE) is used to assess the quality of aligned spatial transcriptome slices in the benchmarking datasets. MSE quantifies the discordance between two sets of 2-dimensional spatial coordinates of tissue spots, typically a query and a reference set. Before calculating the MSE, we translocate and scale the reference coordinates to be between 0 and 10 along both x and y axes, and apply the same translocation and scaling parameters to the query coordinates. This normalization process eliminates the influence stemming from the size of the spatial transcriptome slices on the MSE values, making it easier to cross-reference results from different benchmarking datasets. MSE is defined as the following:

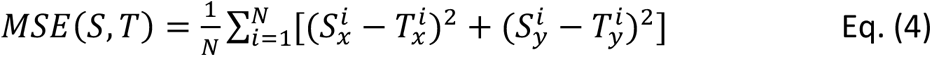

where *N* is the total number of spots, 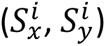 and 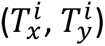 are the normalized coordinates of spot *i* in spatial slices *S* and *T*, respectively.

## Data availability

The H&E staining image of a coronal slice of mouse hindbrain was acquired via Allen Brain Atlas API (31). The Visium spatial transcriptome data from serial sections 1 and 2 (32,33) of a mouse brain sagittal posterior sample were downloaded from 10x Genomics website. The Xenium *in situ* spatial transcriptome data of a mouse brain coronal dissection were downloaded from 10x Genomics website (34). The Visium spatial transcriptome data of a human lymph node sample were obtained from 10x Genomics website (35). The gene counts and tissue image data of the mouse olfactory bulb samples were obtained from the data portal (36) provided by the initial publication of the spatial transcriptome technology (1).

## Author Contributions

Y.B., S.M., and G.S.A. designed the research. P.L., S.M., and Y.B. developed the algorithm. P.L., Y.B., K.X., and S.M. participated in the data analysis. Y.B., P.L., and S.M. wrote the manuscript.

## Acknowledgement

We thank John Sikorski for sourcing and managing GPU cloud computing resources. We also thank Wen Fury for the helpful discussions during the method development and evaluation, and Francesco Randi for the manuscript proofreading.

## Conflict of Interests

P.L., S.M., Y.B., and G.S.A. have filed a patent application relating to the STaCker computational framework. The remaining authors are employees and shareholders of Regeneron Pharmaceuticals, although the manuscript’s subject matter does not have any relationship to any products or services of this corporation.

