## Supplementary Figures for "Image guided construction of a common coordinate framework for spatial transcriptome data"

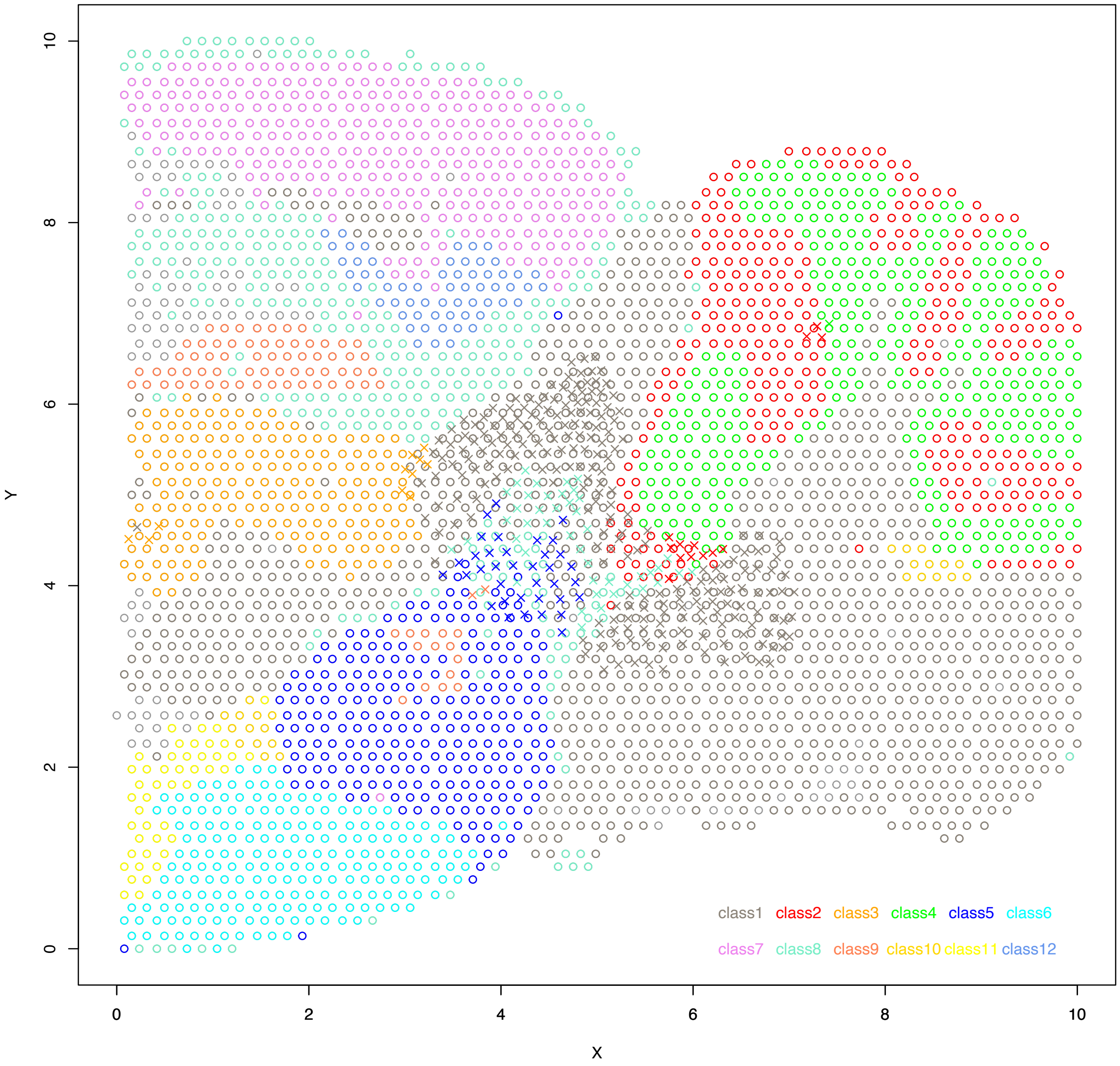


**Supplementary Figure 1**: Spatial consistency of the class labels of the tissue spots in the reference and the STaCker-aligned slide. The presented data is taken from the highly distorted instance in Figure 4 (Simplex noise amplitude = 20, NCC of the deformed image=0.534). The spots in the reference slide (circles) and in the moved slide (crosses) share the same color code of the classes. The displayed spots on the moved slide were aligned by STaCker and have the top 10% largest MSEs relative to their reference counterparts. Note most of the crosses correctly locate in the regions that contain the circles of the same color, indicating their spatial locations are valid regarding their class labels.

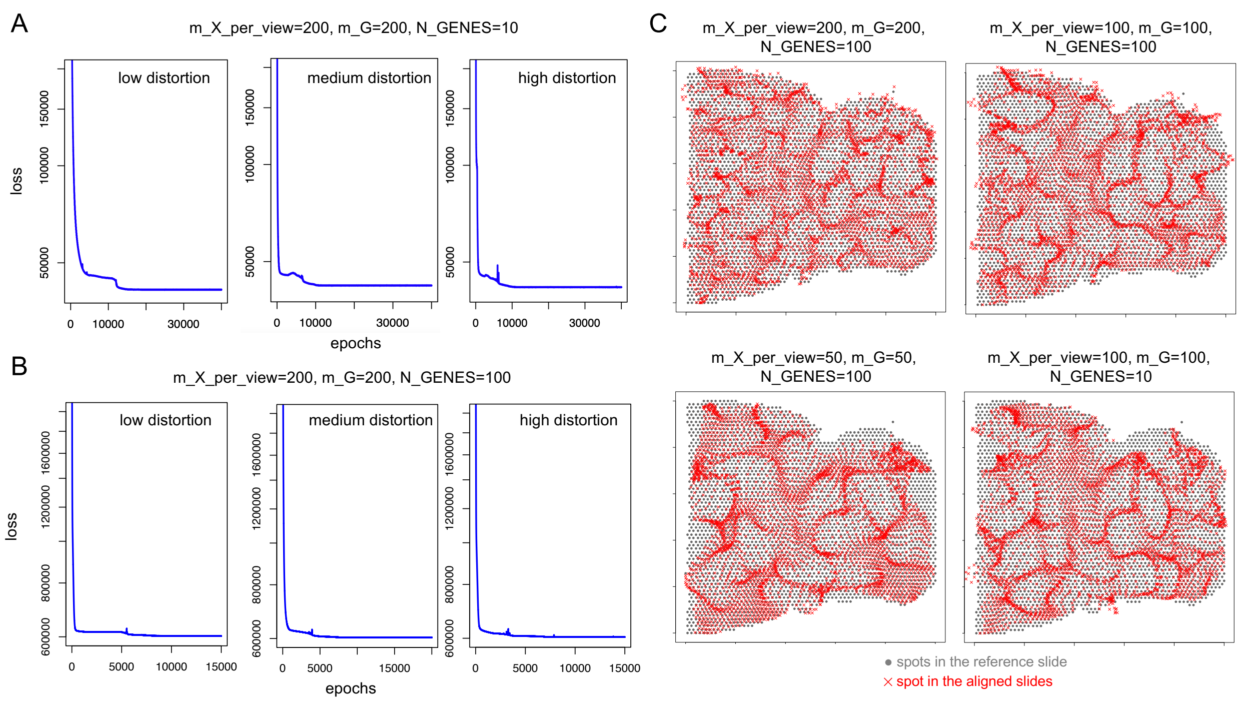


**Supplementary Figure 2**: Exploration of the parameter settings of GPSA in aligning the digitally warped mouse brain slides. A): Trajectories of the loss of GPSA when aligning the moving slides with a low, medium and high distortion in Figure 4. GPSA was executed with the author suggested parameters for Visium platform (m_X_per_view=200, m_G=200, and N_GENES=10). According to the loss trajectory, we ran GPSA for 20,000 epochs or more until the convergence was achieved. B): Trajectories of the loss of GPSA when aligning the same moving slides used in Figure 4 with N_GENES=100 while retaining m_X_per_view=200 and m_G=200. C) The GPSA-aligned spot coordinates (red crosses) are compared to the reference coordinates (grey dots) using the aforementioned and additional parameter settings. In all cases GPSA results in aggregated spatial coordinates.

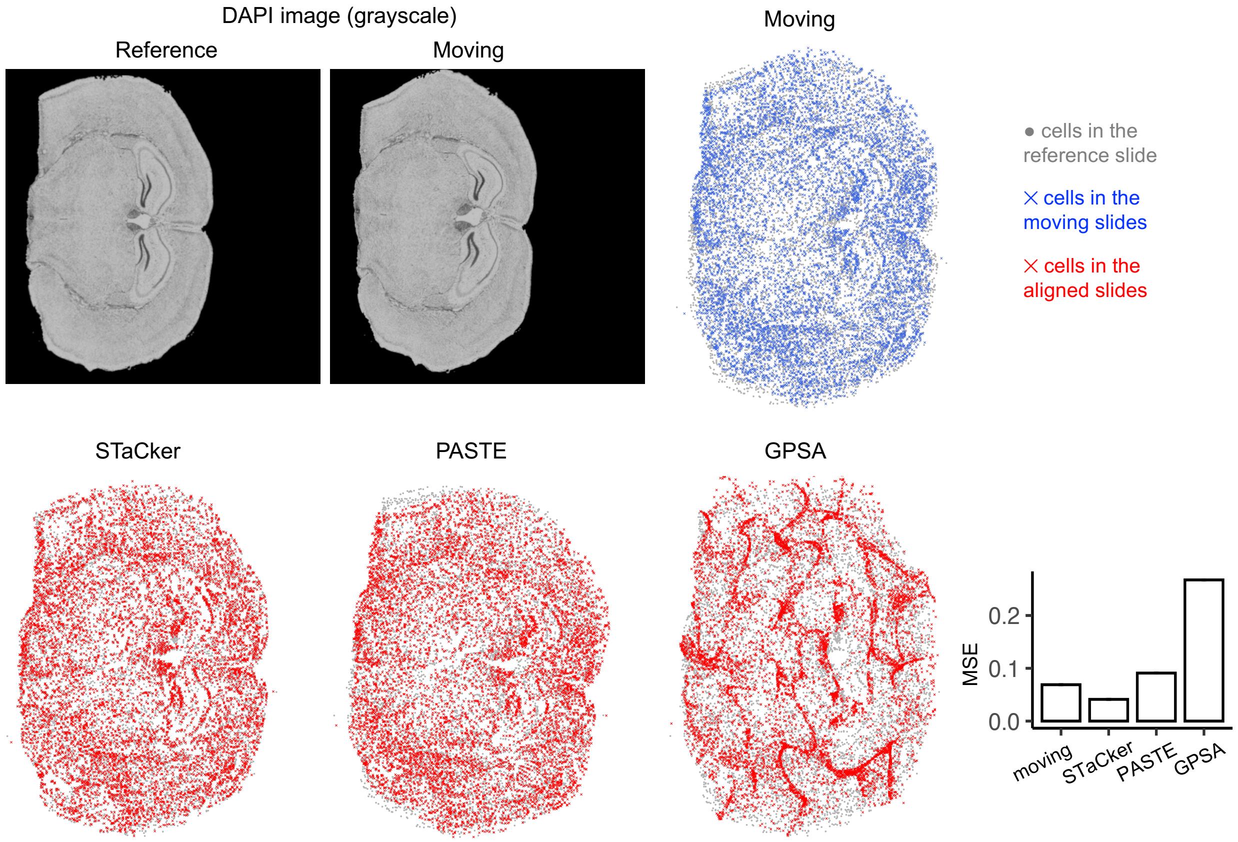


**Supplementary Figure 3**: Evaluation of STaCker in aligning digitally warped *in situ* hybridization (ISH)-based spatial transcriptome slices of mouse brain. The reference is a coronal mouse brain slice containing ~160,000 cells profiled by 10x Genomics Xenium platform. It was digitally warped using Simplex noises (noise amplitude = 15, noise frequency=1, NCC of the deformed image=0.518) to generate the moving slice. The DAPI staining images of the reference and moving slices are shown. As PASTE and GPSA experienced difficulties when aligning all the cells, randomly subsampled 8101 cells were used to quantify and compare the alignments. The coordinates of the 8101 cells in the reference slice (gray dots) are displayed together with their coordinates either in the moving slice (blue crosses) or in the slices aligned by one of the programs (red crosses). The pre-alignment MSE (0.0689) and the post-alignment MSEs (STaCker: 0.0411; PASTE: 0.0909; GPSA: 0.2674) are illustrated in the bar plots. Values from STaCker and GPSA are the average over 10 runs.

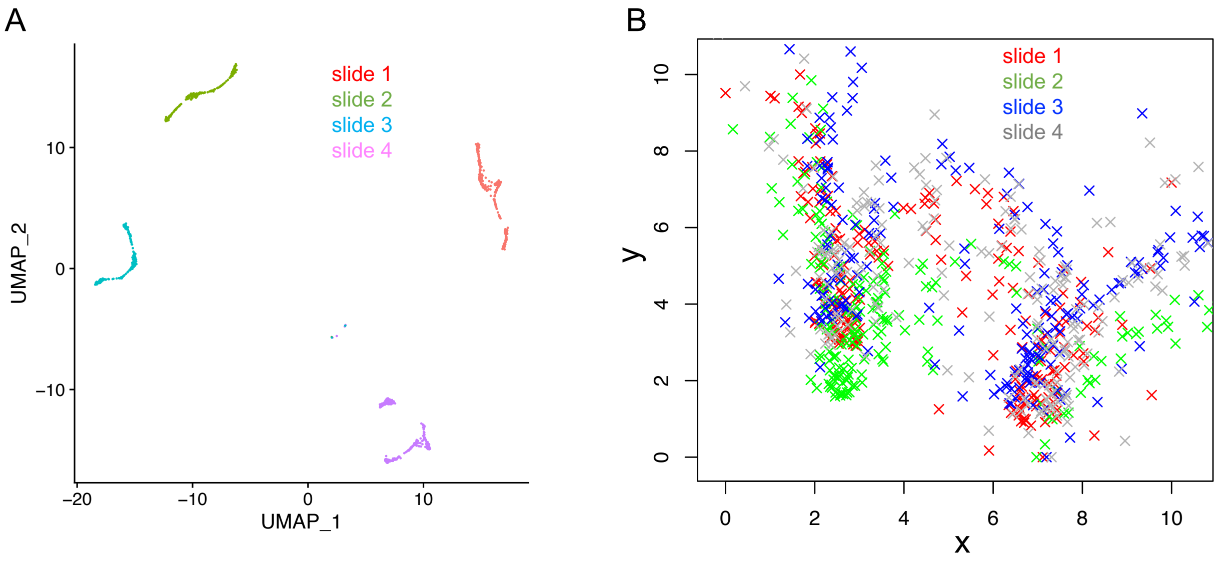


**Supplementary Figure 4**: Batch effect in the mouse olfactory bulb spatial transcriptome dataset. A): The the tissue spots from the four mouse olfactory bulb slides used in Figure 7B were clustered based on their gene expression profiles and visualized in a UMAP. The spots form four slide-specific clusters, indicating there exists a non-negligible batch effect across the slides. B): The tissue spot coordinates from the four slides after the de novo alignment by GPSA. The severe deviation of the coordinates from the expected square grid in Figure 7B reveals the limitation of GPSA. In addition, among the aligned slides themselves, the mutual agreement in the spot coordinates is low (mean pair-wise MSE=24.09), suggesting GPSA is susceptible to the batch effect in the transcriptome data.
